# Human coronary artery tri-culture organ-chip recapitulates anti-inflammatory effect of pulsatile wall strain

**DOI:** 10.1101/2025.09.26.678578

**Authors:** Yu Hou, Georgios Ziakas, Timothy Hopkins, Wen Wang, Hazel R C Screen, Martin M Knight

## Abstract

Inflammation is a precursor to vascular diseases, including atherosclerosis, and is modulated by the local biomechanical environment. There is an urgent need for improved in vitro models, to advance understanding, and to test new therapeutic approaches.. This study describes the development and characterization of a human coronary artery organ-chip model of vascular inflammation, with physiological biomechanical stimulation.

Human coronary artery endothelial cells and smooth muscle cells were cultured on appropriate extracellular matrices in the two adjoining channels of the Chip-S1^®^ from Emulate Inc. Both endothelial and smooth muscle cells demonstrated characteristic phenotypic identity, shown by expression of CD31 and α-SMA respectively. Application of physiological pulsatile tensile strain induced alignment of both cell types, perpendicular to strain direction, as seen *in vivo*. Addition of TNF-α to the vascular channel drove an inflammatory response in both cell types, shown by upregulation of ICAM-1 and P65, and attachment and invasion of circulating THP-1 monocytes. Strain field analysis revealed pressure-dependent spatial variation with 12% strain in the center of the chip, and 5% towards the ends. Pulsatile tensile strain reduced the inflammatory response to TNF-α with a greater localized inflammatory response in areas of lower strain, further replicating *in vivo* behavior.

In conclusion, we present a fully characterized, tri-culture model of the human coronary artery which recapitulates the physiological effects of pulsatile vessel dilation on morphology and localized inflammatory susceptibility. Our model was developed upon a commercially-available, organ-chip platform, allowing for rapid adoption for therapeutic testing, and fundamental discovery science.

## Introduction

Pro-inflammatory signaling within arteries plays a crucial role in cardiovascular diseases such as atherosclerosis; driving the characteristic accumulation of immune cells and buildup of plaque ^1^. The artery is composed of endothelial cells lining the lumen, surrounded by a layer of smooth muscle cells. Arterial inflammation is associated with activation of transcriptional factors, such as Nuclear Factor Kappa-light-chain-enhancer of activated B cells (NF-κB), and the up-regulation of endothelial cell adhesion molecules, such as vascular cell adhesion molecule-1 (VCAM-1) and intercellular adhesion molecule-1 (ICAM-1) ^2, 3^, facilitating monocyte–endothelial cell interaction ^4–6^.

The biomechanical environment of the artery is complex. The endothelium experiences fluid shear stress, due to the flow of blood, and dynamic circumferential strain associated with the pulsatile nature of this flow, whilst smooth muscle cells are exposed solely to the dynamic circumferential strains ^7^. These biomechanical stimuli play a crucial role in artery homeostasis, including the modulation of inflammation. Physiological circumferential strains of 10% have been reported to be atheroprotective ^8–10^. By contrast, areas of the artery where the shear stress and vessel dilation are reduced are more predisposed to inflammation and associated atherosclerosis ^11, 12^. Thus, chronic, localized activation of pro-inflammatory signaling pathways involves a fine interplay between biomechanical and inflammatory stimuli, which regulates vascular health and disease.

Targeting inflammation represents a potential therapeutic strategy for treatment of atherosclerosis and vascular disease. However, both fundamental research and the development of new therapeutics are hampered by the current lack of suitable human *in vitro* models that replicate key features of the artery and associated inflammation. There is currently a reliance on oversimplified in *vitro* models, or non-human *in vivo* models, both of which may fail to accurately recapitulate human physiology and disease, depending on the context of use ^13^. Furthermore, there is now a global push to reduce the use of animals in science and to develop non-animal alternatives. Hence, there is an unmet need for complex *in vitro* models, also known as microphysiological systems, which can be readily translated and adopted to meet both industrial and fundamental research needs ^14^. Organ-on-a-chip (OOAC) systems provide highly controllable, bioengineered microfluidics platforms that recreate the biological, biochemical, and biomechanical environments within an organ so as to replicate key aspects of health and disease ^15^. OOAC systems allow for the incorporation of multiple cell types, and their associated extracellular matrices, to study tissue-specific interactions on a background of physiologically relevant biochemical and biomechanical stimuli ^16–19^. Several groups have applied OOAC systems in studying pathogenesis of cardiovascular diseases and associated drug development ^20–22^. For example, Minkyung Cho et al. engineered a multichannel artery-mimetic chip to study vascular stenosis and inflammation in cocultured layers of endothelial and smooth muscle cells ^23^. Other studies by Mieradilijiang et al. adopted 3D bioprinting approaches to establish a vascular-related tissue model under continuous flow ^24^. However, these and other similar models fail to fully incorporate all aspects necessary to accurately capture arterial physiology or disease. Such factors include the different cell types, their associated ECMs, pro-inflammatory stimuli, circulating immune cells, and physiologically relevant pulsatile flow and/or blood vessel wall dilation. Furthermore, models are often built around bespoke, in-house generated platforms that limit widespread translation and adoption.

In the present study, we report on the development of a novel organ-chip model of the human coronary artery, incorporating human coronary artery endothelial and smooth muscle cells within appropriate ECM, and with physiological pulsatile dilation. We perturb the model by inclusion of pro-inflammatory cytokines and circulating immune cells and demonstrate a clear vascular inflammatory response which is modulated by biomechanical stimuli, as seen *in vivo*. The characterized, non-uniform strain fields measured within the organ-chip enable a single chip to replicate both healthy areas of the artery, and areas that are predisposed to inflammation based on their biomechanical environment, mimicking *in vivo* heterogeneity. By developing this coronary artery model within a commercial organ-chip platform, we provide a robust, scalable model, ideally suited to translation and widespread adoption for therapeutic testing and research.

## Results

### Characterization of human coronary artery organ-chip

Human coronary artery endothelial cells (HCAECs) and human coronary artery smooth muscle cells (HCASMCs) were seeded into the ECM coated bottom channel and top channel of the chip respectively (Fig 2A). After 2 days, the chips were connected to media flow at 30 μl/h. Chips were treated +/- pulsatile tensile strain and +/- 20 ng/ml tumor necrosis factor alpha (TNF-α) for 24 hrs (See methods and Fig 1 for details).

**Figure 1.**
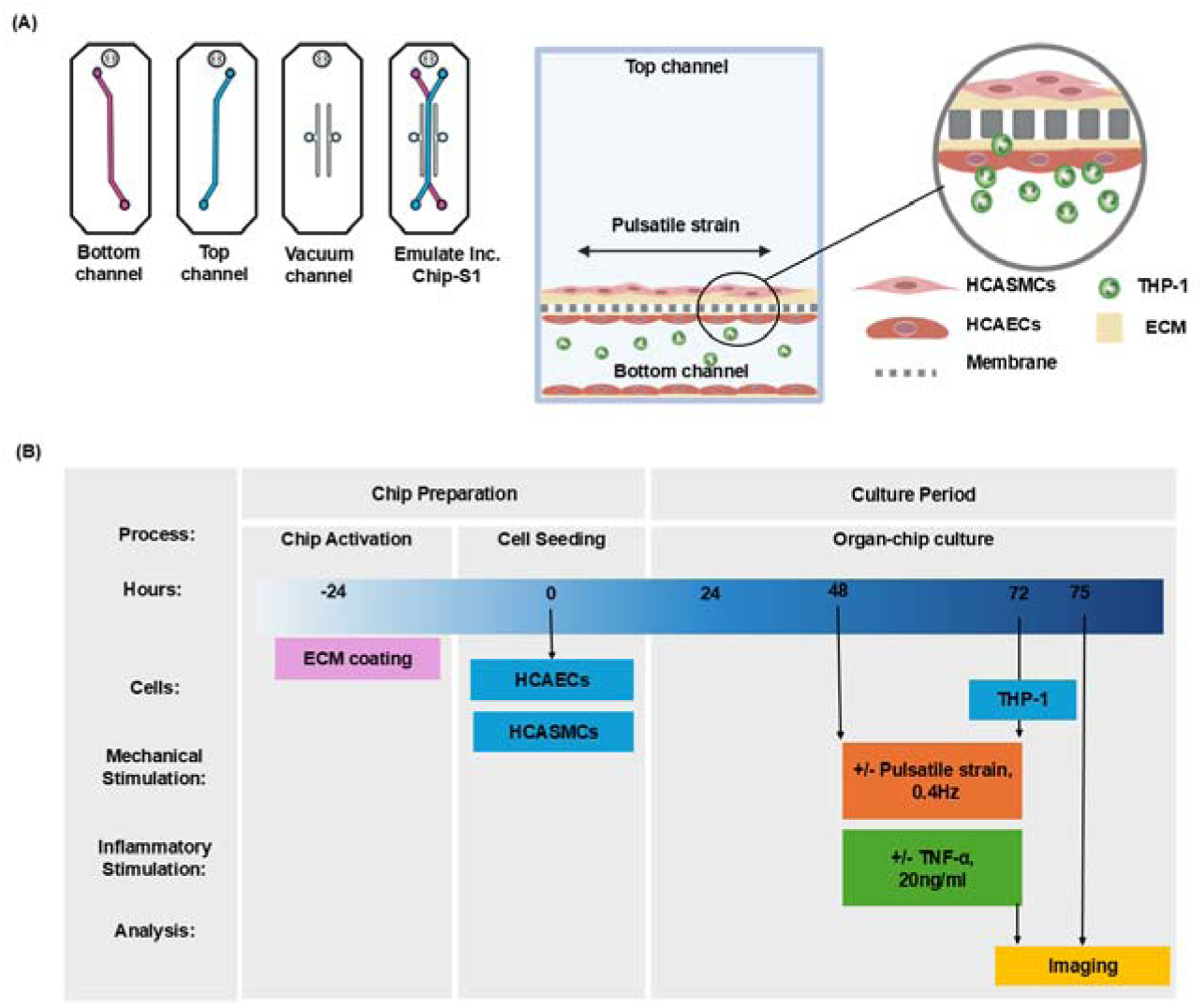
The human coronary artery organ-chip model with controlled pulsatile physiological strain. (A) Exploded schematic of the Chip-S1^®^ from Emulate Inc., showing the two overlapping microfluid channels separated by a porous membrane (7 µm diameter pore size, spaced 40 µm apart in a hexagonal pattern). The top channel (blue) is 1000 µm x 1000 µm whilst the bottom channel (pink) is 1000 µm in width and 200 µm in height. The channels are flanked by two vacuum channels which are used to apply pulsatile strain to the co-culture region. The cross-section shows human coronary artery smooth muscle cells (HCASMCs) cultured in the top channel, and human coronary artery endothelial cells (HCAECs) cultured in the bottom channel both on an extracellular matrix (ECM) coating. Circulating THP-1 monocytes were added in suspension to the culture media in the bottom channel. (B) Schematic highlighting the key steps in the timeline for chip preparation and experimentation showing the application of inflammatory and mechanical stimulation.

**Figure 2.**
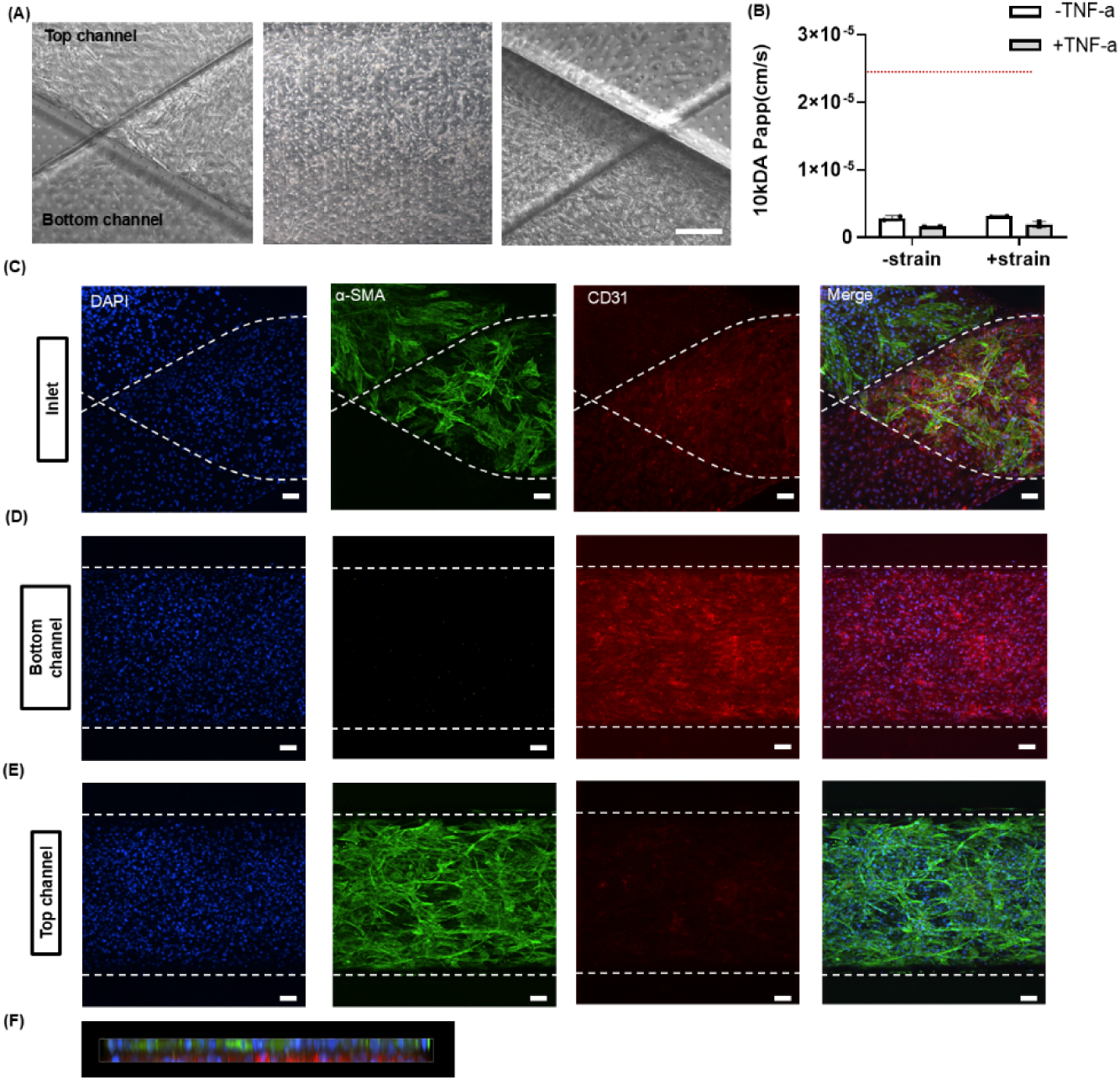
Phenotypic identity within the human coronary artery organ-chip. (A) Brightfield images showing HCAECs evenly seeded within the bottom channel and HCASMCs seeded within the top channel of the Chip S1®. (B) Barrier function assessment of HCAECs and HCASMCs co-cultured chips to determine permeability to 10 kDa dextran under flow. The dotted red line indicates permeability in cell-free organ-chips. Representative images of the inlet region (C) and central regions (D, E) of channels within co-cultured chips labelled for α-SMA (green), CD31 (red) and nuclei (blue). HCAECs and HCASMCs are separated by the membrane (F). Scale bar=100 µm.

The co-culture chips formed a robust barrier impermeable to 10 kDa dextran, with no detectable effect of pulsatile strain and/or TNF-α (Fig 2B) on barrier integrity. By changing the focal plane, separate immunofluorescence microscopy images were obtained of both the top and the bottom channels. This was done at the inlet region (Fig 2C) and in the center of the chip (Fig 2D and E). We demonstrate that the HCAECs in the bottom channel expressed platelet endothelial cell adhesion molecule-1 (PECAM-1)/CD31, whilst in the top channel, the HCASMCs expressed alpha-smooth muscle actin (α-SMA). Thus, we confirm phenotypic identity and distinct localization within the two channels. Hence, the porous membrane effectively maintained spatial separation between the endothelial and smooth muscle channels, as further confirmed by confocal z-x projection (Fig 2F).

We then measured the effects of pulsatile strain, in the presence and absence of TNF-α. Nuclear and cell morphology and orientation, for both HCAEC and HCASMC, were assessed, based on staining with DAPI (Fig 3A and E) and phalloidin (Fig S3). For all conditions, and both cell types, the nuclei were elliptical with aspect ratios greater than 1 (Fig 3B and F). In unstrained chips, HCAECs showed a small increase in aspect ratio in the presence of TNF-α, in the (Fig 3B) with no other effects on aspect ratio or projected area (Fig 3C and G). However, after 24 hrs of pulsatile strain, the nuclei of both the HCAECs and HCASMCs aligned perpendicularly to the direction of the strain (Fig 3C and G). Similarly, the cytoskeleton of both cell types aligned perpendicular to the direction of the strain based on imaging of F-actin (Fig S3).

**Figure 3.**
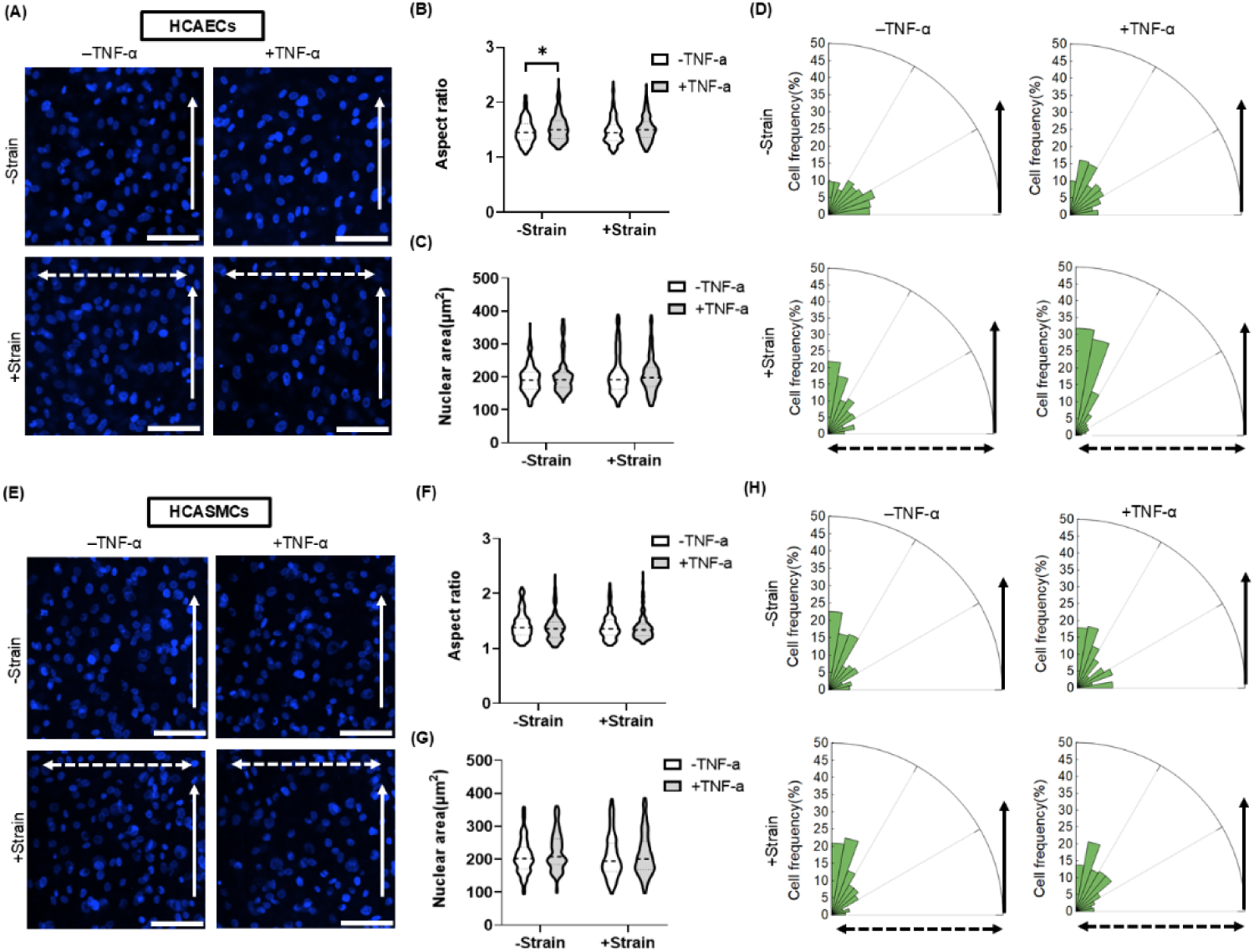
Nuclear morphology and alignment in HCAECs and HCASMCs subjected to unstrained or pulsatile strain in the presence and absence of TNF-α. The co-cultured model was subjected to 24 hrs +/-pulsatile strain, and +/-TNF-α (20ng/ml) treatment. (A, E) Representative confocal images of nuclei stained with DAPI. Scale bar=100 µm. Corresponding data showing (B,F) Aspect ratio, (C,G) Nuclear area (n=200 nuclei) and (D,H) Rose chart of distribution of the nuclear alignment relative to the direction of flow. Arrows indicate flow direction (single-headed) and strain direction (double-headed). Bars represent mean ± SD. Significant differences based on a two-way ANOVA with Bonferroni post hoc and indicated +/- Strain and +/- TNF-α.

### TNF-α induces an inflammatory response within the organ-chip which is suppressed by pulsatile tensile strain

We next investigated how physiological pulsatile strain modulates cytokine induced inflammation in the coronary artery. Off-chip, under static conditions, TNF-α (20 ng/ml) induced a 2-fold increase in ICAM-1 and nuclear p65 expression in HCAECs within 24 hrs (Fig S4A-C). By contrast, TNF-α stimulation of HCASMCs induced a small increase of ICAM-1 but no change in nuclear p65 expression (Fig S4D-F). In the co-culture organ-chip model, HCAECs exposed to TNF-α showed an increase in ICAM-1 and nuclear p65 expression (Fig 4C and D). This pro-inflammatory response in HCAECs was suppressed by pulsatile strain (Fig 4C and D). For the HCASMCs, TNF-α also increased ICAM-1 and nuclear p65 expression (Fig 4G and H). However, these cells were less sensitive to mechanical stimulation with a reduction in nuclear p65 but no change in ICAM-1 (Fig 4G and H). In both cell types, in the absence of TNF-α, strain had no effect on pro-inflammatory markers.

**Figure 4.**
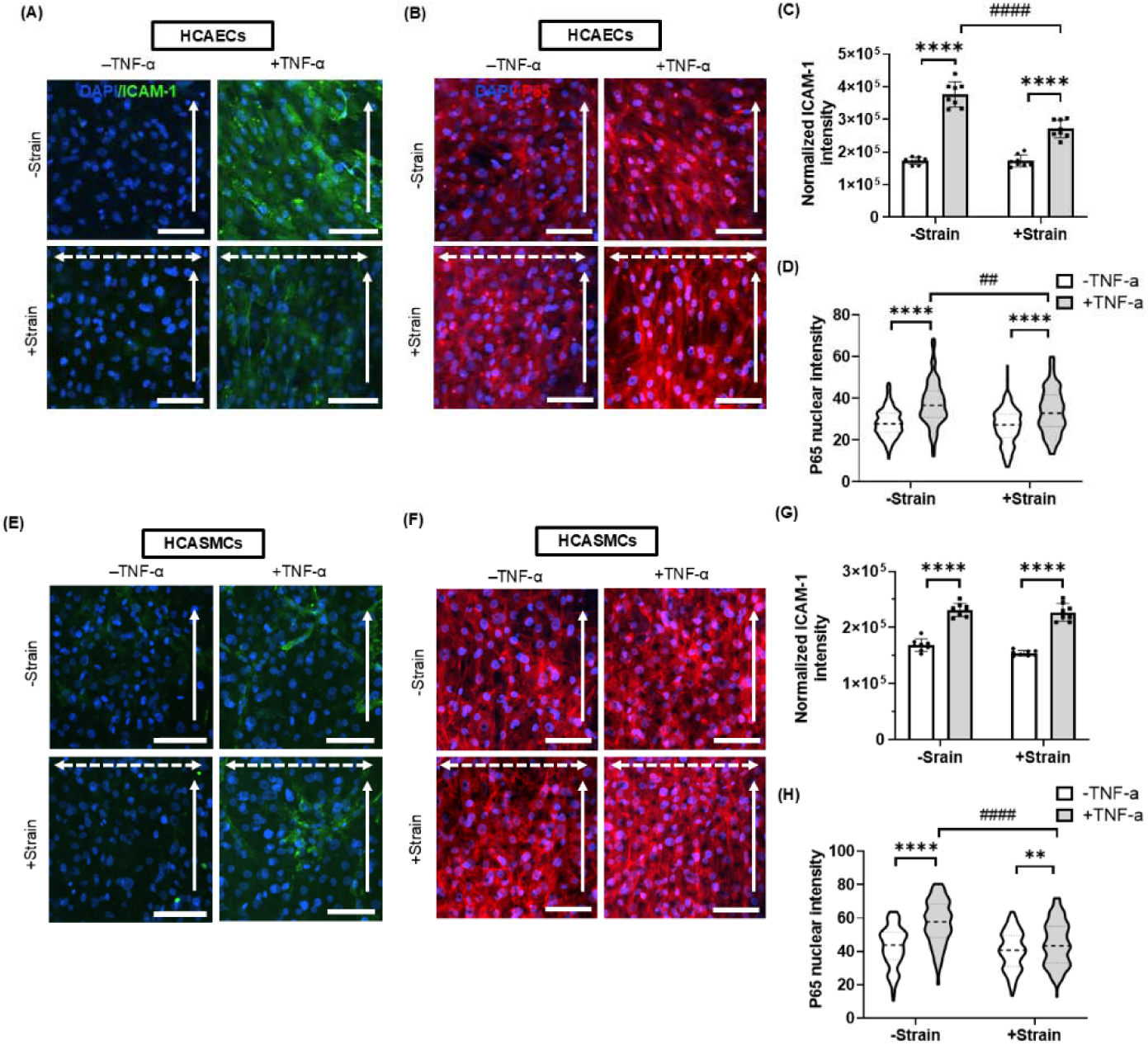
The human coronary artery co-culture model demonstrates a pro-inflammatory response to TNF-α which is suppressed by pulsatile tensile strain. The co-cultured model was subjected to 24 hrs +/-pulsatile strain, and +/-TNF-α (20ng/ml) treatment. Representative confocal images of HCAECs and HCASMCs labelled for ICAM-1 (A and E) and p65 (B and F). Scale bar=100 µm. Corresponding data showing (C and G) normalized ICAM-1 intensity and (D and H) nuclear p65 intensity based on n=8 fields and n=200 cells respectively. Arrows indicate flow direction (single-headed) and strain direction (double-headed). Bars represent mean ± SD. Statistical analysis was based on a two-way ANOVA with Bonferroni post hoc indicated +/- TNF-α (*) and +/- strain (^#^).

These results demonstrate that the human coronary artery organ-chips, with pulsatile strain, recapitulate the reduced susceptibility to inflammation seen *in vivo* in areas of the artery exposed to high laminar shear stress and vessel dilation ^25–27^.

### TNF-α drives monocyte adhesion and migration which is reduced by pulsatile tensile strain within the coronary artery organ-chip

After demonstrating that inflammation was suppressed by pulsatile tensile strain within our co-culture organ-chip, we next extended the model to include circulating immune cells. To quantify monocyte adhesion to endothelial cells and the effect of pulsatile strain, the co-culture organ-chips were maintained for 2 days without flow and then exposed to 24 hrs flow +/- strain, and +/- TNF-α (20 ng/ml). Subsequently, fluorescently labelled THP-1 monocytes were introduced into the endothelial channel and the chip incubated for an additional 2 hrs prior to fixation and microscopy.

Quantification of the images revealed that TNF-α produced a 5-fold increase in the adhesion of fluorescently labelled THP-1 monocytes to the HCAECs in the absence of pulsatile strain (Fig 5A and B). However, the number of adhered monocytes per field of view significantly reduced from 54 ±5.6 (mean ± stdev) in the unstrained chips, to 33 ±4.8 following pulsatile stretch (Fig 5B). Some of the THP-1 monocytes were observed transmigrating through the endothelial channel and into the smooth muscle channel via pores in the organ-chip membrane that separates the two channels. Migration rate was quantified as the ratio of THP-1 monocytes observed in the pores as a percentage of the total adhered monocytes across 8 fields of view. Based on this parameter, 40% of adhered monocytes were found to be in a pore, migrating to the smooth muscle channel in the TNF-α treated chips. However, this was significantly reduced to 25% with the addition of pulsatile strain (Fig 5C). In the absence of TNF-α there were less than 10 monocytes adhered per field of view and none migrating through to the smooth muscle channel. Following treatment with TNF-α, the THP-1 monocytes were clearly visible on the endothelial cells, as shown by confocal imaging of chips, in which HCAECs and HCASMCs were immunolabelled for CD31 and α-SMA respectively (Fig 5D).

**Figure 5.**
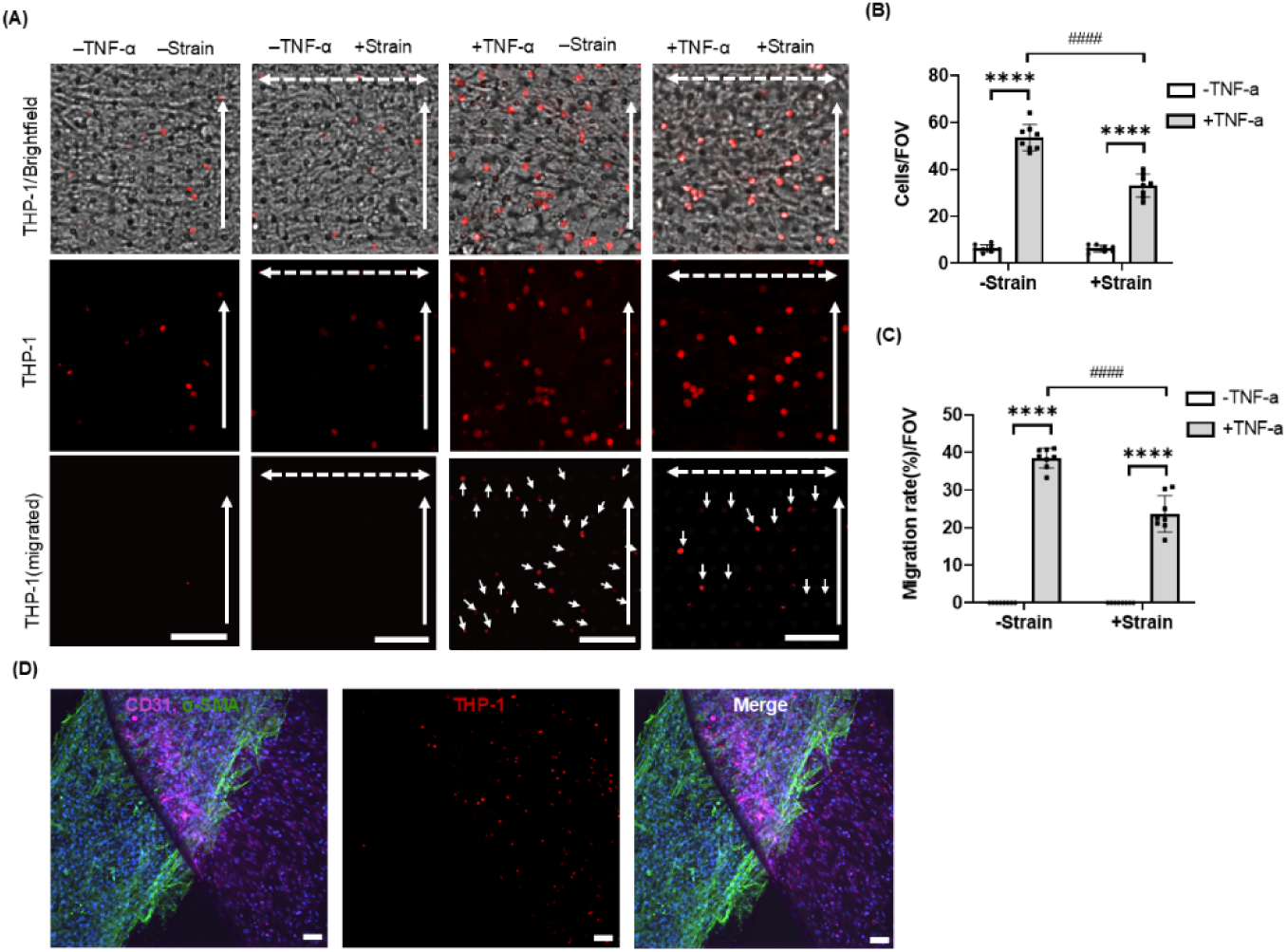
Monocyte adhesion and migration in the inflammatory model of coronary artery organ-chip. The co-cultured model was subjected to 24 hrs +/- pulsatile strain, and +/-TNF-α (20ng/ml) prior to adding THP-1 monocytes. (A) Representative, merge brightfield and confocal images showing THP-1 monocytes adhesion (red) and migration through membrane pores (arrows indicated). Scale bar=100µm. Corresponding quantification of (B) monocyte attachment (n=8 fields), and (C) monocyte migration (n=8 fields). (D) Representative confocal images showing HCASMCs labelled for α-SMA (green) in the top channel, and HCAECs labelled with CD31 (magenta) in the bottom channel, and fluorescent-tagged THP-1 monocytes (red). Nuclei counterstained with DAPI (blue). Arrows indicate flow direction (single-headed) and strain direction (double-headed). Bars represent mean ± SD. Statistical analysis was based on a two-way ANOVA with Bonferroni post hoc indicated +/- TNF-α (*) and +/- strain (^#^).

Collectively, we show that the coronary artery tri-culture organ-chip demonstrates a robust inflammatory response to TNF-α, with increased NF-κB p65 signaling, upregulated ICAM-1 expression, and increased monocyte adhesion and migration. Furthermore, this inflammatory response was suppressed by physiological pulsatile tensile strain, replicating the behavior seen *in vivo*.

### The application of pressure to the Emulate Chip-S1® drives spatial variation in strain parallel and perpendicular to the axis of the channel

Up to this point, all analysis of the effect of pulsatile tensile strain has been conducted in the central region of the microfluidic channels within the Emulate Chip-S1®, away from the inlet and outlet ports. Based on the information from the manufacturer, this region corresponds to a nominal peak strain of 12% at the maximum vacuum pressure of 800mbar. For the final part of this study, we analyzed the 2D strain fields within the entire Emulate Chip-S1® in order to investigate any correlation between localized strain and biological response.

Using brightfield tile scan images (Fig 6A) and the membrane pores as fiducial markers, we successfully measured strains both parallel and perpendicular to the long axis of the channel at different levels of vacuum pressure. This revealed a spatial variation of strain with position along the length of the channel, such that highest strains perpendicular to the channel (ε_width_) were present in the center of the chip (Fig 6B) whilst highest strains parallel to the channel (ε_length_) were found towards the ends of the channels, opposite the ends of the vacuum channels (Fig 6C). The resultant strain magnitude (Fig 6D) and orientation (Fig S8) also varied along the length of the channel.

**Figure 6.**
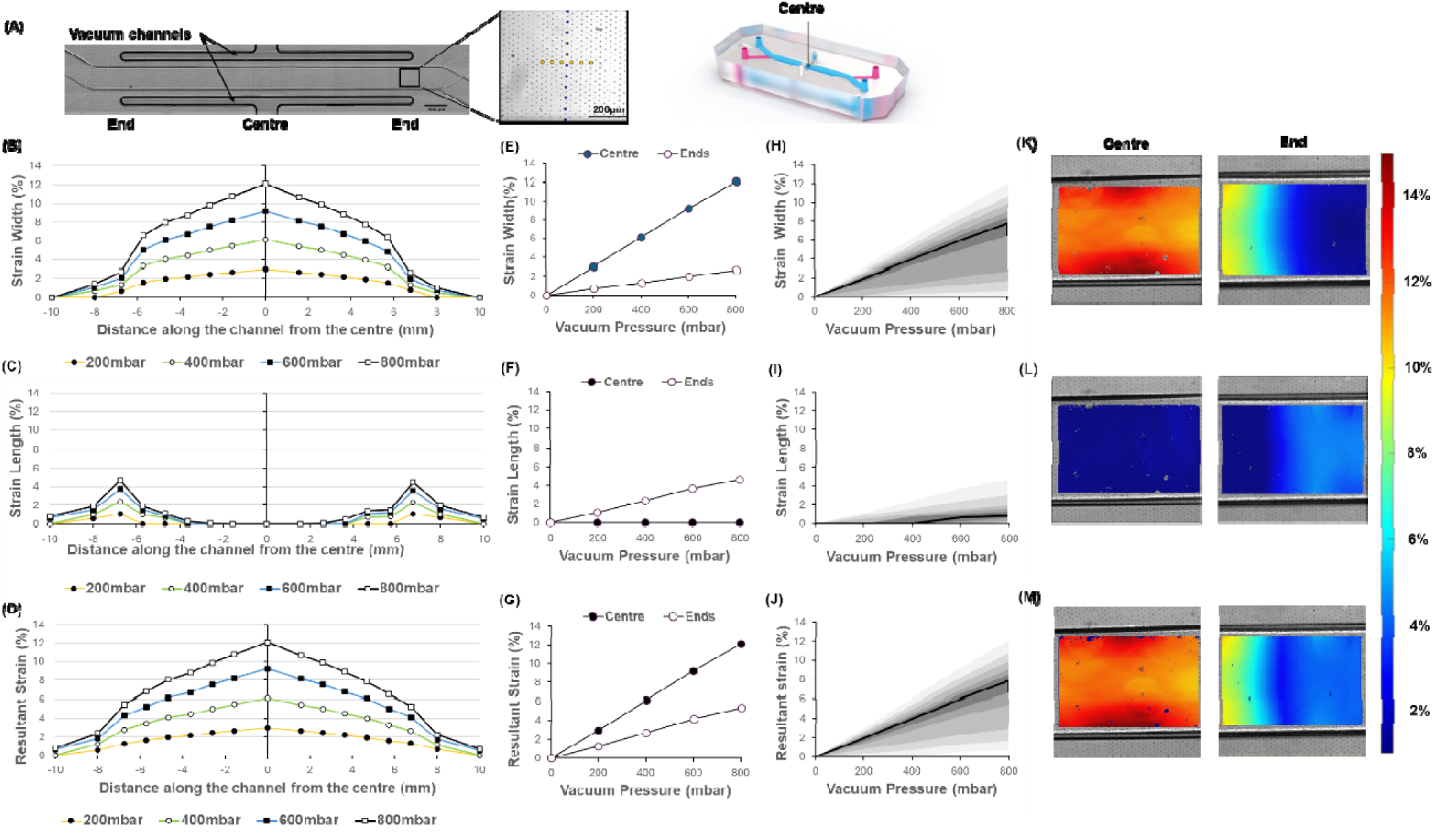
Analysis of strain fields and the effect of vacuum pressure within the Emulate Chip-S1®. (A) Tile scan brightfield image of the Emulate Chip-S1® showing the microfluidic channels separated b the porous membrane and the position of the two vacuum channels either side. Magnified box region shows the hexagonal arrangement of the 7µm diameter pores, and the selection of pores used as fiducial markers. Photo shows the blank chip, in which top and bottom channels have been colored blue and red respectively (right). (B, C, D) Strain profiles along the length of the microfluidic channel at vacuum pressures of 200, 400, 600 and 800 mbar. Strains calculated (B) across the width of the channel, (C) parallel to the length of the channel, and (D) as the resultant strain magnitude. (E, F, G) Corresponding pressure versus strain relationships for strains measured in the centre of the chip and by the end of the vacuum channel as indicated in A. (H, I, J) Corresponding plots showing the median strain (dark line) averaging along the entire length of the channels. Grey shaded regions represent each 0.1 percentile. (K, L, M) Digital image correlation images showing high resolution strain field colourmaps within a single field of view in the centre and end regions (end region corresponds to the righthand side of the chip as shown in A). Color scale indicates the magnitude of strain. See supplementary information for further details (Fig S10).

For all strain measurements at the center of the chip or at the ends (Fig 6E and F), there is an approximately linear relationship between the applied vacuum pressure and the strain magnitude (Fig 6G). Averaging over the entire surface of the membrane, the relationship between applied pressure and distribution of strain values are shown for strains perpendicular (Fig 6H), and parallel to the axis of the channel (Fig 6I) as well as for the resultant strain (Fig 6J). Hence, the median resultant strain at the maximum 800 mbar vacuum pressure is 7.9% with 95% of the surface experiencing a strain between 0.7% and 12.1%.

This analysis confirms that the central region of the channel (1 mm from center), as used for all previous analysis in this study, experienced 10-12% strain, in broad agreement with manufacturers data. However, the end regions, adjacent to the end of the vacuum channels, showed reduced resultant strain magnitude of 5%. We therefore sought to determine whether this variation in biomechanical environment between the center and the ends of the channels was sufficient to induce variation in the anti-inflammatory effect of pulsatile strain within the coronary artery organ-chip model.

### Spatial variation in strain regulates localized endothelial inflammation, recreating both pulsatile high and low strain within a single coronary artery organ-chip

We reanalyzed the previous mechanically stimulated tri-culture organ-chips to quantify the expression of the pro-inflammatory markers in both the central and end regions of the chip where the corresponding resultant strains are approximately 12% and 5% respectively at 800 mbar vacuum pressure (Fig 6G). At the end of the chip where the biomechanical stimulation is reduced, TNF-α induced greater expressions of both ICAM-1 (Fig 7A and C) and p65 (Fig 7B and D), the differences being statistically significant. We also found significantly greater levels of THP-1 monocyte adhesion under lower levels of pulsatile strain at the end of the channels (Fig 7E and F), although there was no difference in migration between the end and the centre (Fig 7G).

**Figure 7.**
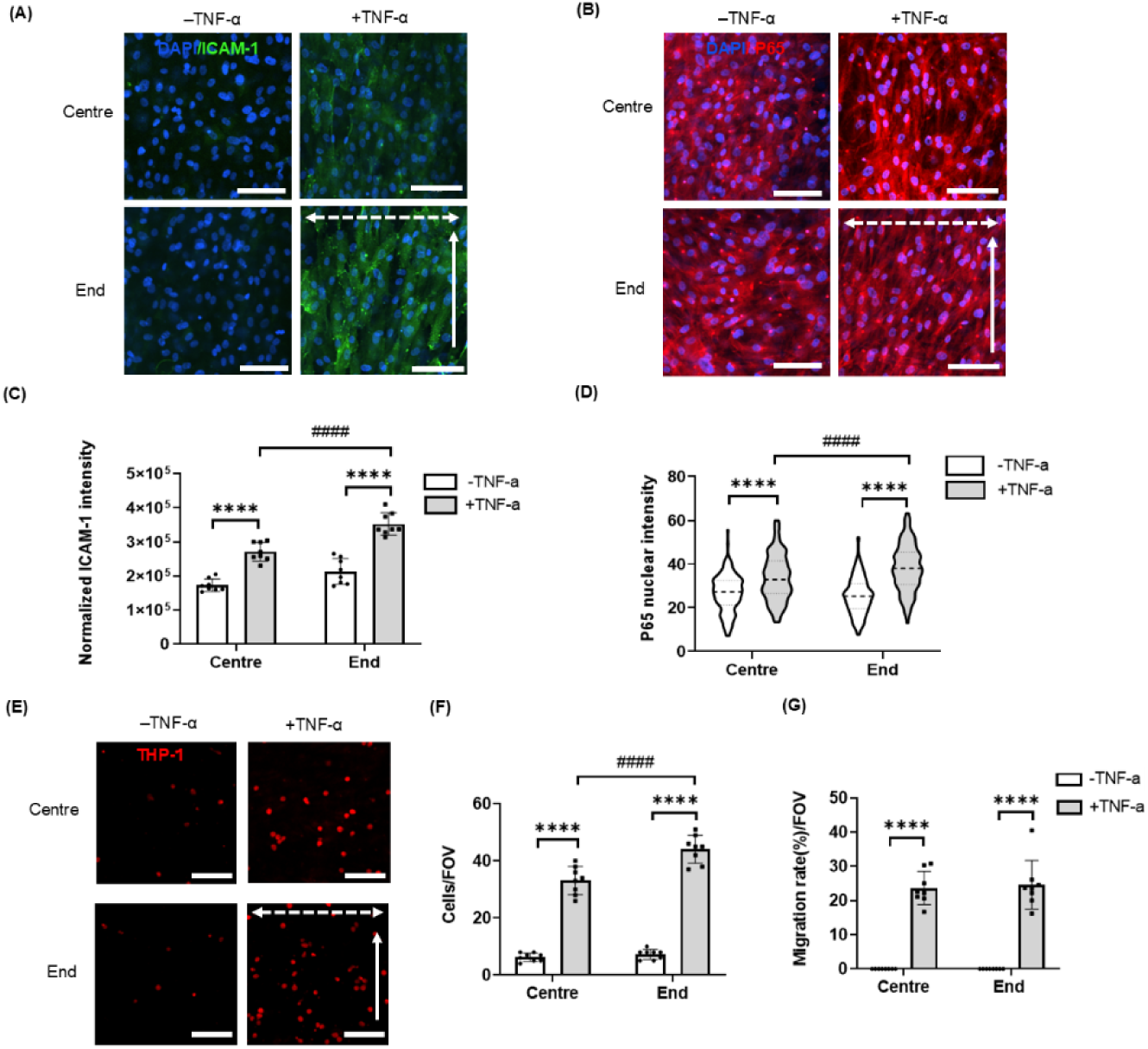
Spatial variation in strain regulates localized endothelial inflammation. The co-cultured model was subjected to 24 hrs +/- pulsatile strain, and +/-TNF-α (20ng/ml) treatment prior to adding THP-1 monocytes. (A) and (B) Representative confocal images of HCAECs labelled for ICAM-1 (green), nuclei (blue) and p65 (red) in Central vs. End regions of Emulate S1 Chip. Scale bar=100 µm. Corresponding data showing (C and D) normalized ICAM-1 intensity and nuclear p65 intensity based on n=8 fields and n=200 cells respectively. (E) Representative confocal images showing THP-1 monocytes adhesion (red). Scale bars=100 µm. Corresponding quantification of (F) monocyte attachment (n=8 fields), and (G) monocyte migration (n=8 fields). Arrows indicate flow direction (single-headed) and strain direction (double-headed). Bars represent mean ± SD. Statistical analysis was based on a two-way ANOVA with Bonferroni post hoc indicated +/- TNF-α (*) and between End and Centre (^#^).

Together these results demonstrate that the spatial variation in tensile strain within different regions of the Emulate Chip-S1® is sufficient to differentially modulate inflammation in response to TNF-α. Hence a single organ chip is able to replicate both the areas of the artery where the biomechanical environment is sufficient to reduce susceptibility to TNF-α, as well as those areas which have a reduced biomechanical environment and are more prone to inflammation.

## Discussion

This study presents a novel organ-chip model of inflammation within the coronary artery, providing a valuable translational bridge for understanding human disease progression and therapeutic responses. A commercially available organ-chip platform was selected in which to create the model in order to maximize adoption without the need for specialist chip fabrication, microfluidics expertise, or in-house technology. Furthermore, the use of a commercial platform maximizes reproducibility and, in the case of the Emulate Chip-S1®, enables highly controlled, physiological mechanical stimuli to mimic pulsatile blood vessel dilation.

The coronary artery is structurally compartmentalized into three functionally integrated layers: tunica intima, which comprises a continuous endothelial monolayer; an internal elastic, tunica media, dominated by smooth muscle cells embedded in elastin-collagen matrices; and the outermost tunica adventitia which anchors the artery to the surrounding tissue ^28^. The organ-chip model presented here replicates this arterial structure. Human coronary artery endothelial cells (HCAECs) formed a monolayer within the bottom channel of chip, positive immunolabelling for the key endothelial marker, CD31 (Fig 2C and D). These cells were co-cultured, via a porous membrane, with human coronary artery smooth muscle cells (HCASMCs) in the top channel, as confirmed by α-SMA labelling (Fig 2C and E).

*In vivo*, within the artery, the endothelial cells are typically elongated and aligned with the direction of flow, and hence perpendicular to the blood vessel wall stretch ^29^. This cell alignment is particularly pronounced in areas of higher fluid shear stress and associated vessel dilation. Similar cell alignment in response to flow alone has also been demonstrated *in vitro* ^30^. This process requires acetylation of microtubules such that loss of the microtubule deacetylase, HDAC6, induced a faster and more robust re-alignment of cells ^31^. In response to tensile strain, many different cell types show changes in cell morphology associated with re-alignment. Generally, continuous static strain induces cell alignment parallel to the stretch direction ^32^, whilst cyclic strain induced cell alignment perpendicular to the stretch direction, as observed in human endothelial and smooth muscle cells under cyclic tensile strain *in vitro* ^33–35^. In the present study we induce physiologically relevant cell alignment of both endothelial and smooth muscle cells, parallel to flow, and perpendicular to the pulsatile tensile strain (Fig 3). Our data suggest that this alignment in the organ-chips is driven by the stretch, since unstrained chips that still had fluid flow, exhibited much reduced cell alignment. However, due to the limitations of the organ-chip platform, the shear stress in the endothelial channel was significantly lower than that likely to occur *in vivo* and was applied in a steady rather than pulsatile manner. Interestingly, in the presence of TNF-α and pulsatile strain, cell nuclei exhibited enhanced perpendicular alignment to the axis of strain (Fig 3), with associated greater reorientation of actin filament (Fig S3).

Dysregulation of arterial function is implicated in the onset and progression of atherosclerosis disease, in which inflammation is common. In human endothelial cells, upregulation of TNF-α increases ICAM-1 surface expression whilst blocking TNF-α can reduce systemic inflammation ^36^. In order to mimic arterial inflammation in the present study, we stimulated cells with TNF-α at 20 ng/ml, as widely used in previous studies to induce pro-atherogenic effects in endothelial cells ^37–39^. TNF-α stimulation within our tri-culture organ-chip resulted in the increased expression of ICAM-1, nuclear p65 translocation, and enhanced monocytes adhesion. TNF-α was introduced only via the endothelial channel, however a pro-inflammatory response was observed in both the endothelial and the smooth muscle cells. Therefore, we performed barrier function analysis to check the permeability to TNF-α. We chose a fluorescently tagged dextran with a molecular weight of 10 kDa since this is slightly less than that of TNF-α at 17 kDa. The cells layer effectively formed a robust barrier to the 10 kDa dextran, indicating that TNF-α applied to endothelial cells would not directly reach the smooth muscle cells (Fig 2B).

Although previous studies have used microfluidic models to examine vascular inflammation and leukocyte recruitment, few, if any, have incorporated smooth muscle cells as well as physiological pulsatile wall strain. In the coronary artery organ-chip presented here, we have characterized the effects of pulsatile tensile strain on cellular organization and cytokine-induced inflammation. Using this model, TNF-α upregulated ICAM-1 expression and p65 NF-κB signaling in both cell types (Fig 4), and increased attachment of monocytes to the surface of the endothelial channel (Fig 5). Thus, this arterial chip effectively mimics the early stages of inflammation and inflammatory cell recruitment as seen *in vivo*. Pulsatile strain suppressed these inflammatory responses, particularly in endothelial cells, recapitulating the *in vivo* situation in which an enhanced inflammatory response is seen in areas with reduced biomechanical stimulation ^40–42^.

## Conclusion

In conclusion, we present a novel artery organ-chip tri-culture model that recapitulates arterial cellular organization and vascular inflammation, with both endothelial and smooth muscle cell components. We demonstrate that pulsatile stretch replicates the previously reported anti-inflammatory effects of physiological pulsatile fluid shear stress ^43^. This enables the model to be created on the commercially available Emulate Chip-S1® to maximize future translation and adoption. The associated strain field heterogeneity enables these organ-chips to effectively mimic areas of the artery exposed to high and low biomechanical environment and to recreate the corresponding differences in the inflammatory susceptibility as seen *in vivo*.

## Materials and Methods

### Cell culture

Human Coronary Artery Endothelial Cells (HCAECs) were purchased from Promocell (C-12221, derived from a single donor, N=1) and maintained in complete Endothelial Cell Growth Medium MV with 5% serum (v/v) (Promocell). This medium was supplied as a kit and consisted of Basal Medium (C-22220) and Supplement Mix (C-22120) with Endothelial Cell Growth Supplement (0.004 mL/ml), Epidermal Growth Factor (recombinant human, 10 ng/ml) and Heparin (90 µg/ml). For stimulation of an inflammatory response, TNF-α (PeproTech), was added to the medium at a final concentration of 20 ng/ml. Hydrocortisone (Promocell, 1 µg/ml) was added only to the medium without TNF-α. Human Coronary Artery Smooth Muscle Cells (HCASMC) were purchased from Promocell (C-12511) and maintained in complete Smooth Muscle Cell Growth Medium 2 with 5% serum (v/v) (Promocell). This medium was supplied as a kit and consisted of Basal Medium (C-22262) and Supplement Mix (C-39262) with Epidermal Growth Factor (recombinant human, 0.5 ng/ml), Basic Fibroblast Growth Factor (recombinant human, 2 ng/ml) and Insulin (recombinant human, 5 μg/ml). Penicillin-streptomycin (P4333, Sigma, 1% v/v) was added to all the medium. THP-1 monocytes (InvivoGen, Toulouse, France) were cultured in Roswell Park Memorial Institute (RPMI) medium (Gibco) supplemented with 10% (v/v) FBS.

### Preparation of the human coronary artery organ-chip

The Chip-S1® (Emulate Inc., Boston, USA) was used (Fig 1A) to generate the coronary artery organ-chip model. This PDMS based chip comprises two channels separated by a thin (50 μm), porous (7 μm pore diameter) membrane. The channels are 1 mm in width, and the channel height is 0.2 mm and 1 mm for the bottom and top channels respectively. Each channel is connected to its own medium reservoir using the Pod™ system (Emulate) such that each cell population can be supplied individually with required media formulations. Two vacuum channels, situated either side of the culture channels, are used to apply pulsatile tensile strain across the membrane (Fig 1A). Prior to cell seeding, the inner surfaces of the channel were activated according to standard protocols (Basic Research Kit Protocol: EP223). 0.5 mg/ml Sulfo-SANPAH (Thermo Fisher Scientific, 22589) was added to each channel and the chips were then treated with UV light for 20 min. Extracellular matrix (ECM), in the form of 33 μg/ml collagen Type I, (C8919, Sigma-Aldrich) was added to the top channel, and a mix of 33 μg/ml collagen Type I and 33 μg/ml fibronectin (C-43060, PromoCell) was added to the bottom channel. Chips were incubated overnight at 37°C. In the final chip configuration, HCASMCs were cultured in the top channel and HCAECs in the bottom channel (see schematic in Fig 1A). As shown in Fig 1B, following the ECM coating of the two channels, chips were washed with pre-warmed culture medium and HCAECs seeded to the bottom channel. Chips were inverted for 3 hrs to encourage attachment to the membrane, then righted. HCASMCs were then seeded to the top channel and allowed to attach for 3 hrs prior to gentle washing.

After a further 48 hrs, chips were connected to the automated culture module (ZÖE; Emulate) and cultured with their respective media flowing at a rate of 30 μl/h. Both cell types were subjected to 24 hrs pulsatile tensile strain at a frequency of 0.4 Hz by application of 800 mbar to the vacuum channels. Based on manufacturer’s instructions, this was expected to generate a strain in the central region of the chip of approximately 10-12%. The associated pulsatile distortion of the microfluidic channels is likely to induce some degree of pulsatile fluid flow, particularly in the bottom channel, although this has not been quantified. Control chips were left unstrained throughout. Inflammatory stimulation was provided by addition of a single dose of 20 ng/ml TNF-α to the endothelial channel.

### Barrier function assessment

To assess barrier function in co-culture chips, an experiment was carried out in which the culture media in the bottom channel contained a fluorescent tracer dye (0.1mg/ml FITC-conjugated 10 kDa dextran (Sigma-Aldrich)). After 24 hrs of flow +/- strain, and +/- TNF-α, chips were disconnected from the Zoë and 100 μl of media was collected from each of the bottom and top channel outlet reservoirs. Fluorescence intensity was measured at 355 nm and 485 nm, and the concentration of dye was determined for the dosing channel (bottom channel) and receiving channel (top channel) using a standard curve. This data was used to assess the apparent permeability (Papp) according to the following calculation:

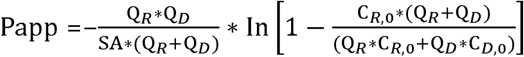

where Papp is the apparent permeability in units of cm/s, SA is the surface area of the co-culture channel (0.17 cm^2^), Q_R_ and Q_D_ are the fluid flow rates in the receiving and dosing channels, respectively, in units of cm^3^/s, and C_R,0_ and C_D,0_ are the recovered concentrations in the receiving and dosing channels respectively (Emulate).

### Immunofluorescence staining and confocal microscopy

Cells were fixed with freshly defrosted 4% paraformaldehyde (PFA, 10min, room temperature) and permeabilized with 0.5% Triton-X/PBS (5min, room temperature). Subsequently, the samples were blocked for 1 hr at room temperature with 20% Fetal Bovine Serum (FBS, F7524, Sigma), incubated overnight at 4°C with primary antibodies, washed in 0.1%BSA/PBS (3×10min), and incubated with secondary fluorescent-conjugated antibodies (Table 1). Cell nuclei were counterstained with DAPI (D9542, Sigma). Confocal z-stack imaging was performed using a Nikon CSU-W1 SoRa Spinning Disk Confocal laser scanning microscope with a 20x/0.75/NA objective. Confocal z-series were obtained consisting of 15 sections with a nominal z-spacing of 0.6 µm. Maximum intensity projections were created and the images analyzed using ImageJ (NIH). ICAM-1 integrated intensities were measured using ImageJ. Normalized ICAM-1 integrated intensities were determined by dividing integrated intensity by cell number for each field of view. The quantification of nuclear NF-κB p65 was performed based on methods previously described (Fig. S1) ^44^. The quantification of nucleus orientation and aspect ratio was performed using PAT-GEOM, a free software package based on ImageJ software (Fig. S2) ^45^.

**Table 1.** Antibodies used for immunofluorescence staining (IF) and western blotting (WB)

| Target | Feature | Cat no./species | Company | Dilution |
| --- | --- | --- | --- | --- |
| CD31 | ECs marker | ab9498/rabbit | Abcam | 1:1000 (IF) |
| α-SMA | SMCs marker | ab7817/mouse | Abcam | 1:1000 (IF) |
| ICAM-1 (CD54) | Inflammatory marker | ab2213/ mouse | Abcam | 1:1000 (IF) |
| Phalloidin | F-actin | sc-363797 | Santa Cruz | 1:1000 (IF) |
| NF-κB (P65) | Inflammatory marker | ab32536/ rabbit | Abcam | 1:1000 (IF) |
| Alexa Fluor 488 | Secondary antibody | A21202/ mouse | Invitrogen | 1:1000 (IF) |
| Alexa Fluor 555 | Secondary antibody | A32794/rabbit | Invitrogen | 1:1000 (IF) |
| Alexa Fluor 647 | Secondary antibody | A21245/rabbit | Invitrogen | 1:1000 (IF) |

### THP-1 monocyte adhesion and migration assay

THP-1 monocytes were labelled with 2 μM cell tracker dye (CMRA orange, Thermo Fisher Scientific) for 15min in serum-free medium at 37°C. Cells were then centrifuged at 300 ×g for 5min to form a pellet, washed with serum-free medium then resuspended to a concentration of 2×10^6^ cells/ml. Fluorescently labelled monocytes were perfused into the bottom channel of the chip then incubated under static conditions at 37 °C for 2 hrs. The channel was then flushed with Endothelial Cell Growth Medium to remove unattached cells, and chips were imaged to assess monocyte adhesion and migration using confocal microscopy. The quantification of monocyte adhesion and migration was performed by counting labelled monocytes with ImageJ software (Fig. S5).

### Analysis of strain fields within the Emulate Chip-S1®

A Fluigent® push-pull controller was connected to the vacuum channels of the Emulate Chip-S1® which was mounted on the Nikon CSU-W1 SoRa Spinning Disk Confocal. Using a 10x/0.3/NA air immersion objective, brightfield images of the porous membrane separating the two microfluidic channels were obtained at 0, 200, 400, 600 and 800 mbar of negative pressure applied to the vacuum channels. Tile scanning was used to create a series of images which were automatically stitched together to make a single image covering the full length of the chip at each pressure (Fig. 6A). The pores were then used as fiducial markers for analysis of strains at different positions along the length of the channel. For each position, the strains were calculated parallel and perpendicular to the long axis of the channel, termed ε_length_ and ε_width_ respectively (Fig S6). The resultant strain magnitude and angle of orientation relative to the long axis of the channels were then calculated (Fig S8).

### Digital Image Correlation (DIC) for 2D Strain Analysis

DIC was performed using Ncorr, an open source 2D DIC software ^46^. Emulate Chip-S1® were prepared as previously described. However, to provide sufficient texture for the correlation algorithm, chondrocytes were isolated as previously described ^47^ and seeded on the membrane in the top channel. For the DIC analysis the following parameters were used: Subset Radius: 25 pixels, Subset Spacing:10 pixels, Strain Radius: 25 pixels, Iteration Cutoff: 50 iterations, Step Analysis, RG-DIC Subset Truncation and Strain Subset Truncation: disabled. Resulting pseudocolour maps were created showing the magnitude for each strain parameter within the field of view (Fig S10).

### Statistics and data analysis

Statistical analysis was performed using GraphPad Prism version 8 for Windows (GraphPad Software, San Diego, California USA). Statistical analysis and n values for each experiment are outlined in the figure legends with significant differences indicated at p<0.05 (* or ^#^), p<0.01 (** or ^##^), p<0.001 (*** or ^###^) and p<0.0001 (**** or ^####^).

## 1. Supplementary Information

See the supplementary material for additional data and methods.

## 2. Acknowledgements

YH was supported via a China Scholarship Council PhD studentship in collaboration with Queen Mary University of London. GZ was funded on a PhD studentship from Queen Mary University of London. TH was funded on a Versus Arthritis Fellowship working on organ-chip models of inflammation (Versus Arthritis Foundation Fellowship #22876). We are grateful to support of the Queen Mary in vitro Models facilities which have been funded via an equipment grant from NC3Rs (www.cpm.qmul.ac.uk/facilities).

## 3. Author contribution

YH, GZ, TH and MK conceived and planned the studies. TH provided practical input on organ-chip use, inflammatory stimulation, and monocyte culture. YH conducted all the experiments involving the arterial cells. GZ conducted the strain field analysis. All authors were involved in data analysis and drafting the manuscript.

## 5. Ethics approval

Ethics approval is not required.

## 6. Data availability

The data that support the findings of this study are available from the corresponding authors upon reasonable request.

