## Supplemental materials for "Human coronary artery tri-culture organ-chip recapitulates anti-inflammatory effect of pulsatile wall strain"

**Manuscript category**: Research Article

**Keywords**: organ-on-a-chip, artery, inflammation, microphysiological systems, mechanobiology, biomechanics, atherosclerosis

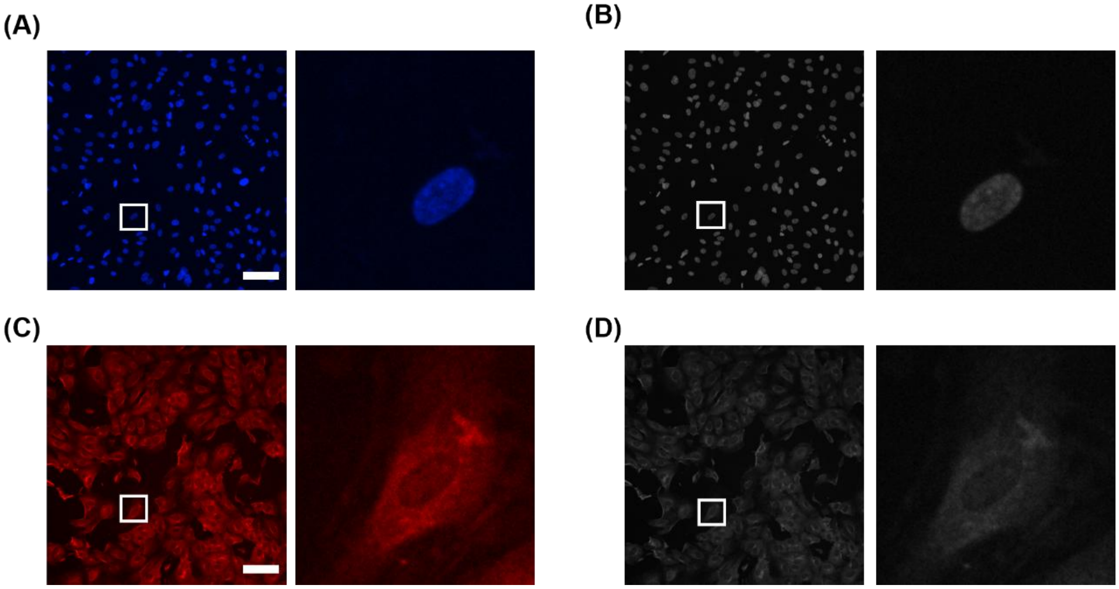


**Fig. S1 Method for quantitation of p65 nuclear intensity.** Representative confocal images of (A) DAPI and (C) P65 from the same field of view. Scale bar=100 μm. Thresholded image (B) were generated from the DAPI stained image (A). Thresholded image (D) were done for p65 stained image (C). Following particle analysis, a data file of numerical values and drawing with numbered nuclei is generated, which the nucleus ROIs was applied to the thresholded p65 image to quantify nuclear p65 intensity.

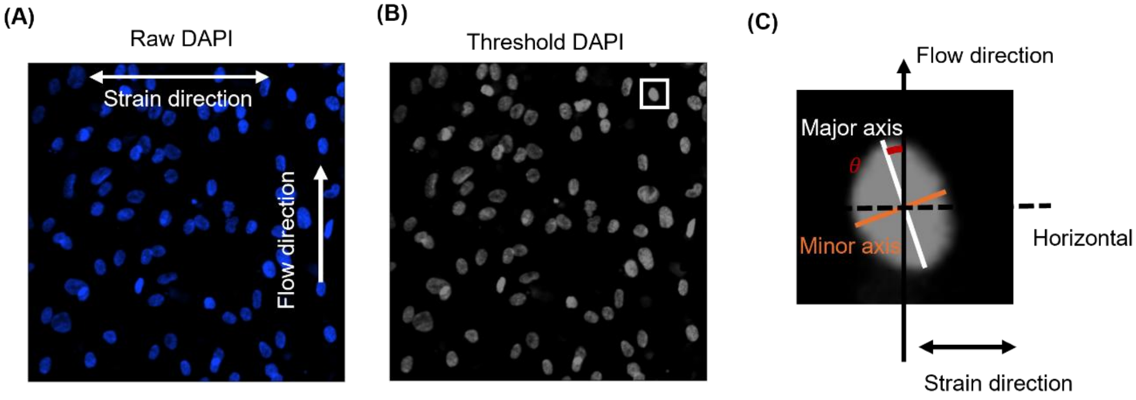


**Fig. S2 Method of analyzing nuclear orientation.** (A) Each raw confocal maximum projection image of nuclei stained with DAPI was converted using ImageJ, to (B) an 8-bit thresholded image. This was used for automated image analysis such that each nuclei was numbered and matched to a best fit elipse. (C) The aspect ratio (major/minor) was calculated for each nuclear elipse and nuclear orientation was determined as the angle along the major axis relative to the direction or flow (or long axis of the channel in no flow conditions).

### Supplementary Information

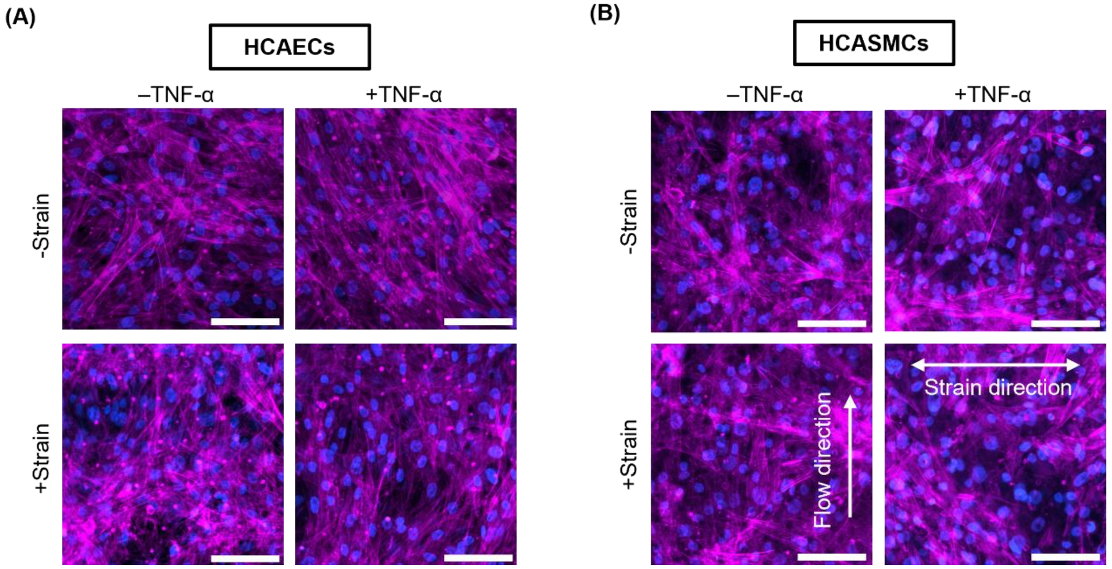


**Fig. S3 Actin alignment in HCAECs and HCASMCs subjected to unstrained or pulsatile strain in the presence and absence of TNF-α.** Co-cultured chips were subjected to pulsatile strain for 24 hrs and during treatment with +/-TNF-α (20ng/ml), unstrained was used as a control. Representative confocal images for HCAECs (A) and HCASMCs (B) of actin (purple) and nuclei stained with DAPI (blue). Scale bar=100 µm.

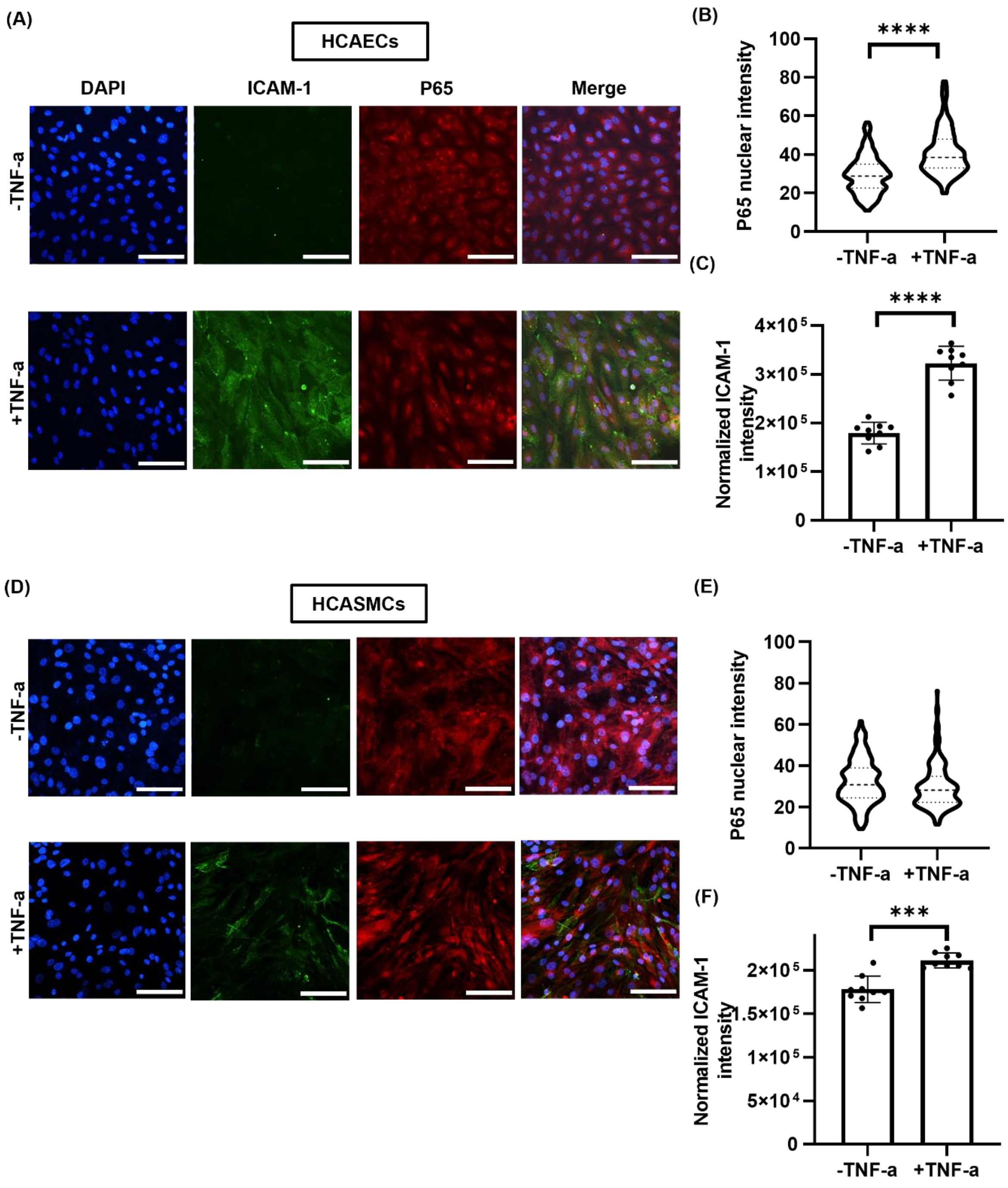


**Fig. S4 Tumor necrosis factor (TNF)-α incubation leads to up-regulation of inflammation in HCAECs and HCASMCs.** Cells were treated with +/- TNF-α (20 ng/ml) for 24 hrs. Representative confocal images of (A) HCAECs and (D) HCASMCs. Cells were labelled for nuclei (DAPI, blue), ICAM-1 (green) and p65 (red). Scale bar=100 μm. Corresponding analysis showing, (B and E) p65 nuclear intensities (n=200 cells), and (C and F) normalized ICAM-1 intensities (n=9 fields). Bars represent mean ± SD. Statistical analysis was based on Mann-Whitney U test for (B) and (E), Student t test for (C) and (F).

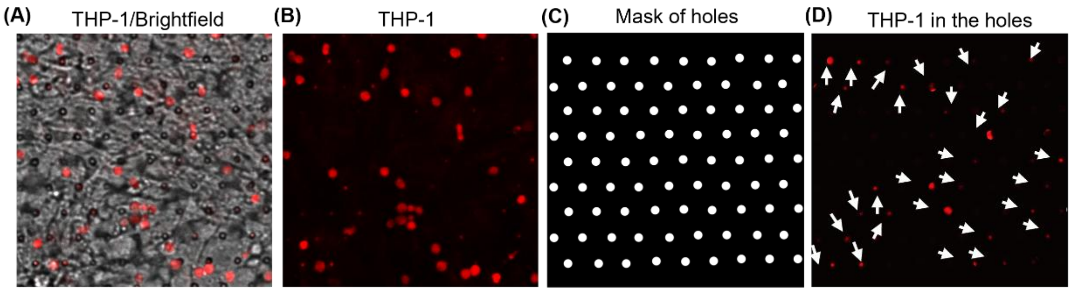


**Fig. S5 Method for identifying THP-1 monocyte migration in the organ-chip.** Representative confocal image of fluorescently-tagged THP-1 monocytes (A) with and (B) without overlay brightfield image showing the porous membrane. (C) Corresponding mask showing the position of pores in the membrane based on the brightfield image. (D) Merged image of (B) and (C), to indicated those pores in which a monocyte was present (arrows).

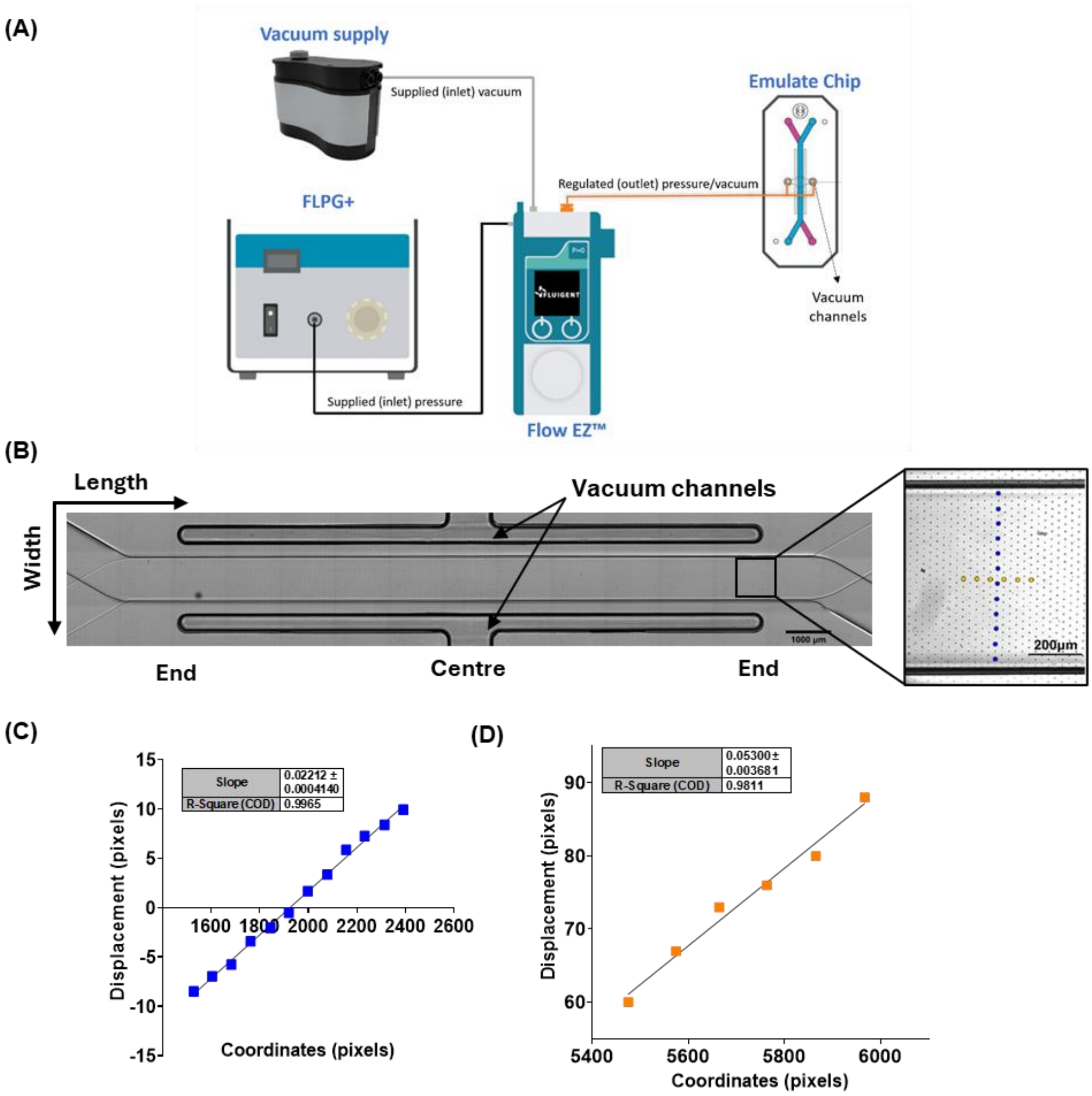


**Fig. S6 Method for calculating 2D strain fields along the length of the channel based on analysis using fiducial markers in Chip-S1®.** (A) Schematic of the Fluigent pressure control system connected to Emulate Chip-S1 vacuum channels. (B) Tile scan brightfield image of the Emulate Chip-S1® showing the microfluidic channels separated by the porous membrane and the position of the two vacuum channels either side. Magnified box region shows the hexagonal arrangement of the 7µm diameter pores with the selection of pores that are used as fiducial markers (blue and orange dots) for displacement analysis. The blue pores are used to track the displacement across the width of the channel, while the orange pores provide displacement along the length. Corresponding graphs showing displacement versus original position, (C) across the width, and (D) along the length of the channel. Linear models were fitted to the data such that the gradient indicates the strain.

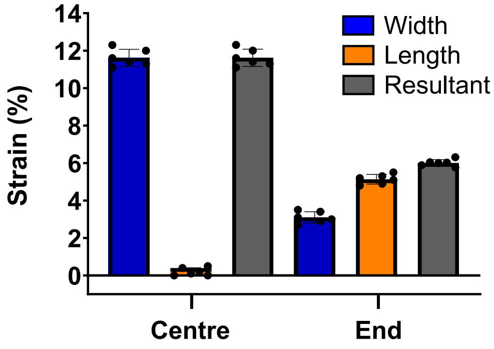


**Fig. S7 Reproducibility of tensile strain within in Emulate Chip-S1®.** Data shows strains measured in both the centre and ends of the channel, perpendicular (blue) parallel (orange) to the long axis of the channel, defined as $\varepsilon_{width}$ and $\varepsilon_{length}$ respectively. In addition, the resultant strain (grey) is also indicated. Strain was measured 3 times at each location in two different chips, n=6 measurements (dots). Bars represent mean ± standard deviation.

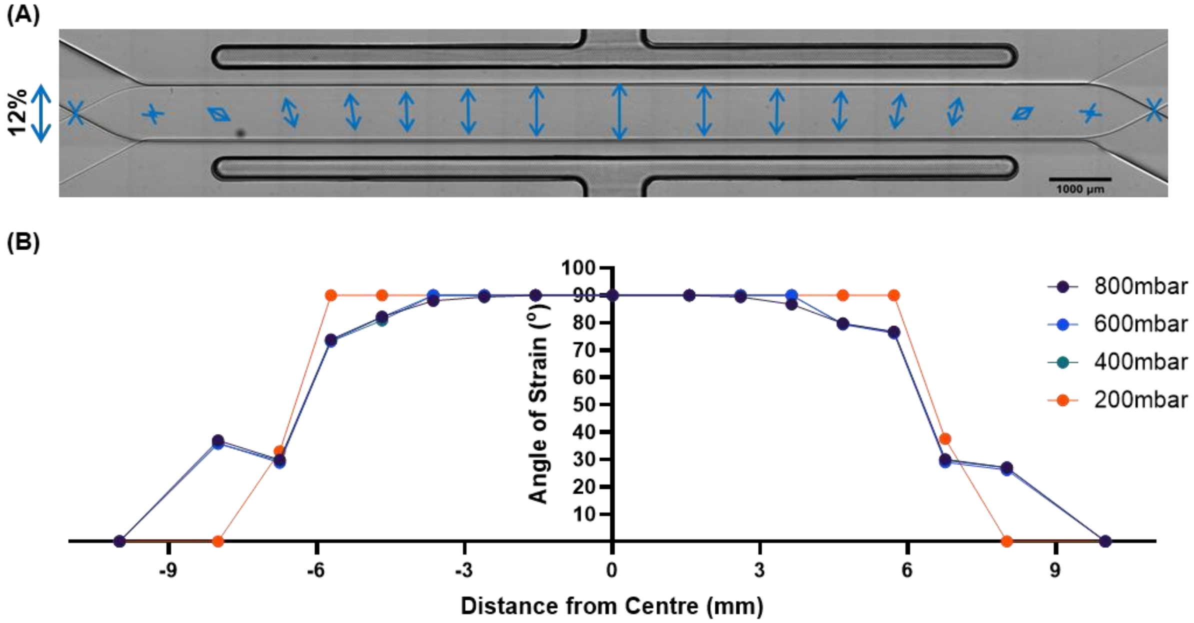


**Fig. S8 Angle of resultant strain at different positions along the length of the channel in the Emulate Chip-S1®.** (A) Tile scan brightfield image of the Emulate Chip-S1® with blue arrows indicating the magnitude and angle of the resultant stain at multiple positions along the channel length. For scale the arrow on the left indicates a resultant strain of 12% at an angle of 90°. Values correspond to an 800mbar vacuum pressure. (B) Corresponding quantification of the angle of strain relative to the length of the channel, measured at increasing vacuum pressures (200, 400, 600 and 800 mbar). Data show that the angle of strain remains close to 90° (perpendicular to the length of the channel) along the central region and decreases towards the channel ends.

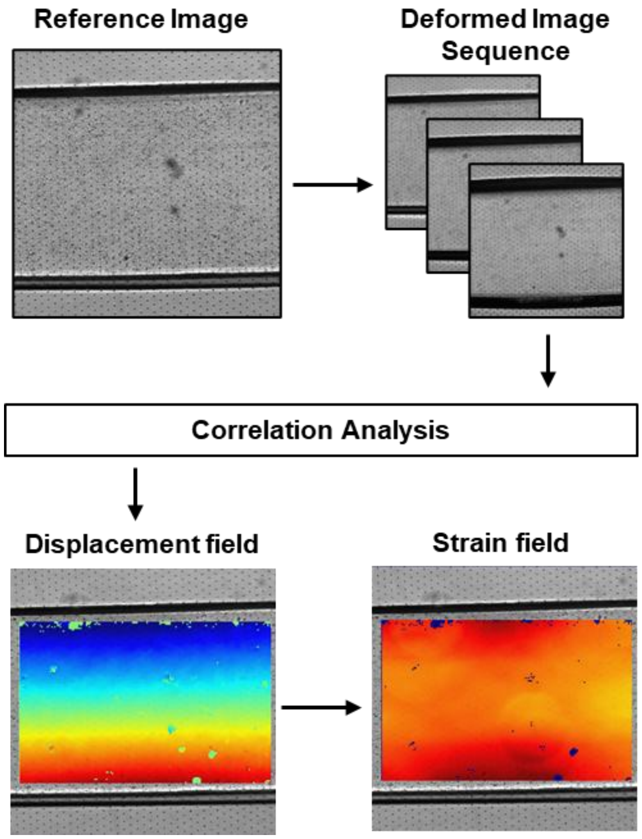


**Fig. S9 Workflow for applying Digital Image Correlation to analyse 2D strain in the Emulate Chip-S1®.** A reference image of an unstrained chip is first acquired, followed by a sequence of images captured under increasing vacuum pressure (200, 400, 600 and 800 mbar). Correlation analysis is carried out using Ncorr (MATLAB). This divides the image into subsets and compares the reference and deformed images to calculate local displacements, generating a displacement field (bottom left). The displacement data are then used to compute the 2D strain field (bottom right), enabling full-field mapping of strain distribution within the region of interest.

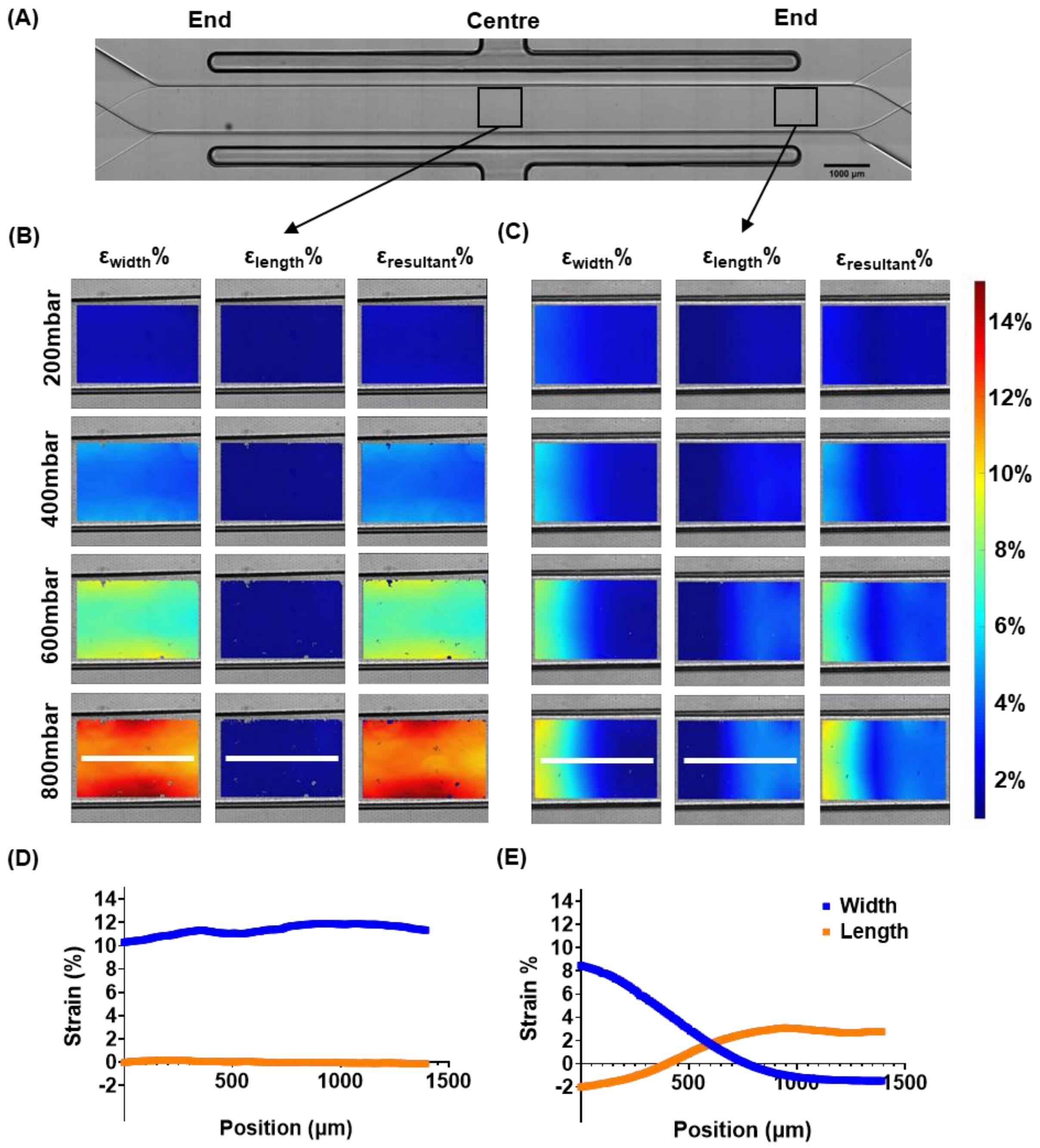


**Fig. S10 Digital Image Correlation reveals spatial variation in strain in the Emulate Chip-S1®.** (A) Tile scan brightfield image of the Emulate Chip-S1® showcasing the different regions analysed with DIC. Corresponding strain maps, (B) at the centre, and (C) at the end of the channel, showing the strain across the width of the channel ($\varepsilon_{width}$%), along the length ($\varepsilon_{length}$%), and the resultant strain ($r_{resultant}$%). Data presented at 200, 400, 600, and 800 mbar of vacuum pressure. Colour bars indicate strain magnitude from 0% (dark blue) to 14% (dark red). Corresponding strain profiles measured as indicated by white lines on the strain maps at 800 mbar with data for $\varepsilon_{width}$ and $\varepsilon_{length}$, (D) at the centre, and (E) at the end of the chip.
